# Language Model Embedding Classifiers Enable Identification of Multiple Sclerosis-Associated BCRs and Repertoires

**DOI:** 10.64898/2026.07.07.735316

**Authors:** Graham C. Peet, Gregory P. Owens, Jeffrey L. Bennett, Arjun Krishnan, Wendy B. Macklin

**Affiliations:** Neuroscience Program, University of Colorado Anschutz Medical Campus, Aurora, CO, USA; Department of Cell and Developmental Biology, University of Colorado Anschutz Medical Campus, Aurora, CO, USA; Department of Neurology, University of Colorado Anschutz Medical Campus, Aurora, CO, USA; Department of Ophthalmology, University of Colorado Anschutz Medical Campus, Aurora, CO, USA; Program in Immunology, University of Colorado Anschutz Medical Campus, Aurora, CO, USA; Department of Biomedical Informatics, University of Colorado Anschutz Medical Campus, Aurora, CO, USA

## Abstract

Multiple sclerosis (MS) is a chronic inflammatory demyelinating disease. It affects over 2 million people worldwide but has historically been challenging to diagnose, categorize, and treat. MS has an autoimmune component that involves the production of B-cell receptors (BCRs) and immunoglobulins (Igs) that are associated with disease pathophysiology. We reanalyzed all publicly available RNA sequencing data from MS patients and extracted over 11 million BCR immunoglobulin heavy chain (IGH) sequences obtained from a variety of tissue sources. We developed a new decoder-only BCR DNA embedding model that outperforms state-of-the-art BCR and general-purpose protein language models on sequence embedding tasks. We then trained a language model classifier capable of identifying BCR sequences associated with MS. Using low-dimensional representations of whole repertoire embeddings combined with sequence-disease predictions, we can distinguish MS patient repertoires from healthy, infectious disease, or other autoimmune disease repertoires. Our models also successfully rank known MS-associated myelin-binding IgG sequences relative to controls. These findings provide a methodological foundation for BCR-based MS detection and could facilitate identification and study of disease-associated antibodies from blood.

**Significance:** Autoimmune diseases are difficult to diagnose and treat. Circulating adaptive immune cells contain accessible information about a patient’s present and past immune experiences. To better use this information in the context of multiple sclerosis, we extracted a comprehensive dataset of B-cell receptor sequences from archival data, developed a new DNA foundation model to embed these sequences, and applied machine learning methods to classify sequence disease association and patient disease state. These findings will advance our ability to model and predict multiple sclerosis and other diseases.

## Introduction

Like many diseases with an autoimmune component, multiple sclerosis (MS) has historically been difficult to study and diagnose. Even with advanced imaging and modern diagnostic criteria, a meaningful proportion of MS cases represent misdiagnosis, and delayed diagnosis is common (1, 2). Although the underlying pathological process in MS is still not understood, experimental evidence indicates diverse involvement of the adaptive immune compartment, and B cell targeting therapies have proven effective at reducing inflammatory activity and slowing disease progression (3). In the last decade, targeted T and B cell receptor repertoire sequencing has enabled insight into the diverse and informative changes that occur in adaptive immune receptors in MS (4–6). Yet, a diagnostic adaptive immune signature for MS remains elusive.

A number of promising technologies exist or are in development to better capture the immune changes characteristic of MS. Machine learning optimization methods have been used to identify biochemical properties of fragmented CSF BCR sequences associated with relapsing-remitting MS (RRMS) with a reasonable degree of accuracy (7). Boosting tree models applied to microarray data have shown promise in distinguishing MS from non-autoimmune neurological diseases and among MS subtypes (8). More recently, very accurate models have been developed to integrate protein language model BCR embeddings to accurately classify complex autoimmune conditions like type I diabetes and systemic lupus erythematosus (SLE) (9). These increasingly sophisticated tools indicate that disease-specific signals in MS can be identified in BCRs and show that powerful language model technology can be applied to extracting BCR disease-specific signals.

Language models have shown strong capability to classify adaptive immune receptors and patient diseases. The structure of Igs consists of constant and variable regions, with the constant regions providing the structural scaffold of the molecule. The variable regions that mediate binding to diverse epitopes are produced through somatic recombination of gene segment arrays and subsequent mutation that occurs following B cell activation and affinity maturation. Each antibody is generated by a clonal B cell lineage that may be as small as a single cell; individuals contain approximately 10^9^ circulating B cells (10). This extremely large search space makes characterizing disease-relevant antibodies and their targets difficult. Due to their moderate sequence length and high complexity, BCR sequences are well-suited to analysis with transformer-based language models (11, 12). Such language models have recently sparked breakthroughs in related areas such as protein folding and antibody design, and we believe the problem of identifying an MS-associated BCR signature is ripe for this approach (13, 14).

Given that their unique BCR gene sequence determines specificity and that Igs have a role in MS pathology, understanding whether BCRs across the MS disease states have common sequence features may help to identify and classify B cell clones most relevant to MS pathology. Toward this goal, we performed an exhaustive reanalysis of BCR sequences in archived data from MS patients and controls, extracting millions of BCR IGH sequences not previously analyzed. We then developed novel language processing tools to produce high-quality unsupervised embeddings and supervised sequence-disease association predictions. These tools create more informative embeddings, which allows us to classify MS-associated sequences with greater accuracy and produce equal or better patient classification relative to existing methods. Importantly, this work could enhance our ability to accurately diagnose MS in patients at early stages of the disease, when novel therapeutics may be particularly effective in delaying overall disease progression (15).

## Results

### A comprehensive retrospective dataset of B and T cell receptors from MS patients and controls

We accessed all human sequence runs available in December of 2024 in the NCBI sequence read archive (SRA), in response to the search query “multiple sclerosis”. We manually deduplicated submissions and removed runs such as microRNA sequencing that could not contain adaptive immune sequences. Due to extremely sparse metadata, many sequence runs had to be manually annotated using information from the original publications. TRUST4, a repertoire assembly tool for unselected RNA sequencing, was then used to extract and assemble BCR sequences in the filtered files (16). We additionally accessed data from the OPIG OAS repository of unpaired BCR sequences and re-assembled several in-house datasets containing MS and neuromyelitis optica (NMO) BCR sequences (4, 17, 18). Among SRA-derived sequences, we captured approximately 800 distinct patients, 63 percent of whom were female (**Fig. 1A**). Patients ranged in age from 12 to 92 with similar mean ages between MS and retrieved control samples (**Fig. 1B, C**). Both published studies and the number of patients sampled increased in availability around the year 2019 as RNA sequencing studies became more prevalent (**Fig. 1D**). In total, we were able to assemble 11,563,296 total BCRs, including 2,818,768 high-quality sequences with full VDJ gene segment assembly (**Fig. 1E**). Analysis of this comprehensive dataset allowed us to identify IGH V gene usage enriched in MS patient blood samples relative to control patients included in the studies accessed from SRA. Additionally, sequences enriched in MS patient CSF samples relative to MS patient blood samples were identified, including V genes such as IGHV4-39 and IGHV2-26 previously identified as associated with MS (**Fig. 1F, G**) (19, 20). To address the question of shared IGH CDR3 clones between MS patients (“public clones”), we computed the enrichment of public clones between patients within the same study and between patients across studies relative to the expected empirical rate of public clones based on CDR3 amino acid length and sequence sampling depth. We found that public clones are significantly enriched within studies, indicating frequent technical overestimation of these events (**Fig. 1H**). We also compared the mean CDR3 region length in MS patients to control patients, finding a detectable increase in length commonly associated with autoimmune conditions or chronic immune stimulation in MS patient CSF but not blood samples (**Fig. 1I**) (21). The comprehensive dataset investigated here is larger and contains more varied patients than previously available resources, and our findings related to V gene usage, public clones, and CDR3 length demonstrate the validity and utility of the dataset.

**Fig. 1:**
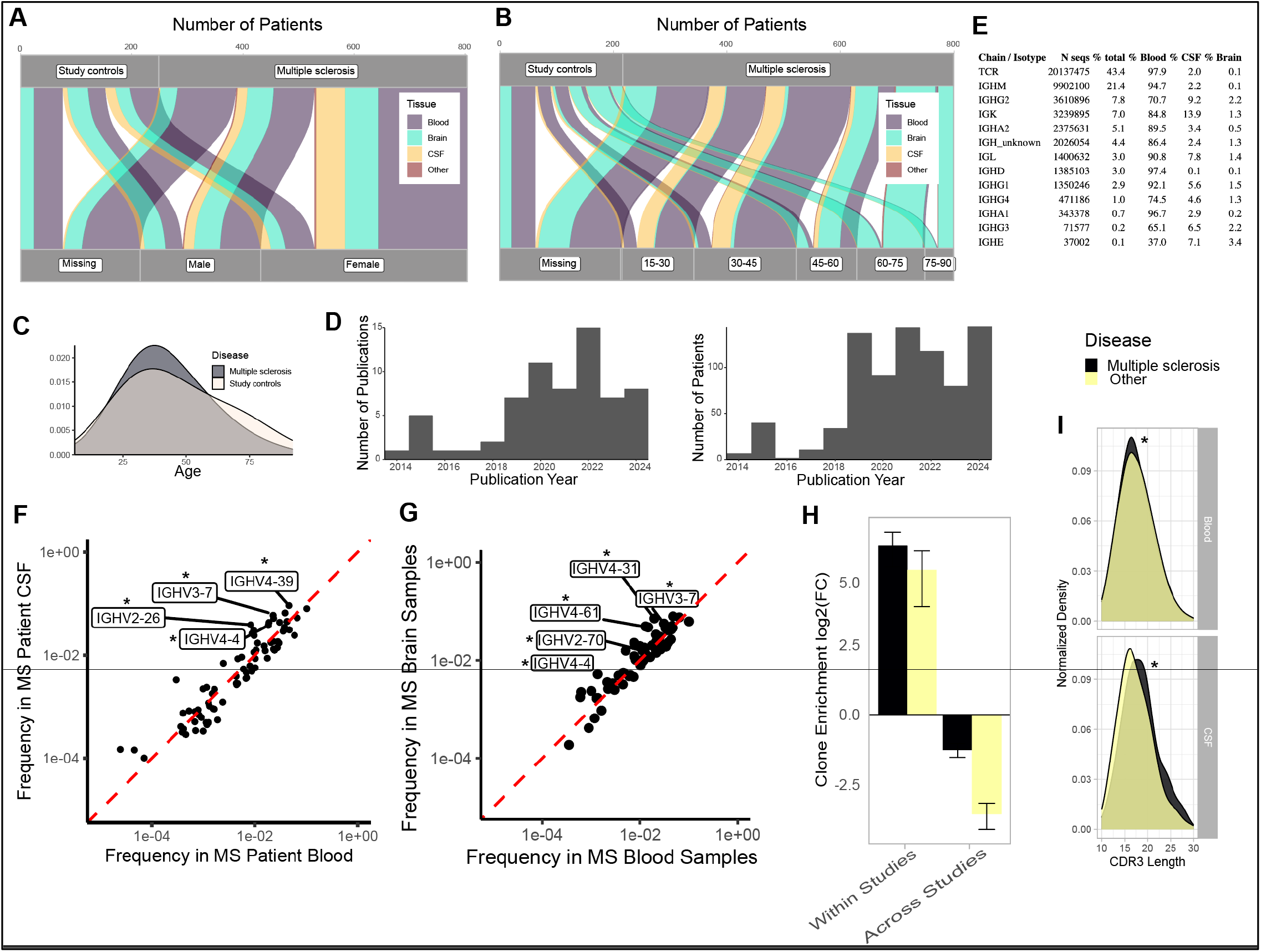
A comprehensive retrospective dataset of B and T cell receptors from MS patients and controls *A)* Relationship between TRUST4 assembled MS patients and associated controls from studies in NCBI SRA by sex and tissue source. *B)* Relationship between TRUST4 assembled MS patients and associated controls from studies in SRA by age and tissue source. *C)* Distribution of MS patient and associated study control patient ages. *D)* Temporal distribution of publications and patients retrieved from NCBI SRA. *E)* Table of retrieved adaptive immune sequences from MS samples and associated controls by chain/isotype, and tissue source. *F)* Enrichment of IGH V genes in MS patient CSF samples relative to MS patient blood samples from SRA studies. * = p < 0.01 *G)* Enrichment of IGH V genes in MS patient brain samples relative to MS patient blood samples. * = p < 0.01. Wilcoxon rank-sum test for FDR adjustment. *H)* Frequency of IGH CDR3 clones between MS patients or control patients within the same study compared to the frequency of IGH CDR3 clones between MS patients sampled from different studies, relative to the expected empirical public clone rate across all sequences assembled. *I)* CDR3 (junction) length in MS patient blood and CSF compared to control blood and CSF. * = p < 0.001. Student’s T test with FDR adjustment.

### A BCR-specific DNA foundation model and classifier for MS-associated BCRs

Because of the strong performance of general-purpose and tailored language models in many adaptive immune sequence embedding tasks, we next used public non-MS data to pretrain a DNA language model tailored to BCRs for the purpose of better understanding our MS BCR dataset. Because small BCR-specific protein language models can outperform even larger general-purpose protein language models at BCR embedding tasks and because a DNA rather than a protein language model can capture all protein level information while also capturing genetic patterns not evident at the protein level, we trained a new unsupervised foundation model based on downscaled variants of the Mistral-7b architecture (BCR-Mistral) (**Fig. 2A**) (22, 23). We assessed how model size impacted predictive training on three variants with approximately 100 million, 300 million, and 500 million parameters, finding that the largest model substantially outperformed the other two models on validation set data for both the entire BCR IGH sequence and the CDR3 alone (**Fig. 2B**). We used BCR IGH data from OPIG OAS exclusively to pretrain the model, and preselected heterogeneous sequences based on nucleotide 7-mer counts to improve training speed and generalizability. To benchmark model performance, we prepared a dataset of antigen-adapted (<95% V gene identity) productive full-length sequences from SARS-CoV-2, myasthenia gravis, and SLE patients, as well as MS patient CSF samples. We benchmarked performance by measuring the relative distances in embedding space between positive (same disease, different patient) and negative (different disease, different patient) sequences sampled at random and by computing the cosine similarity between sequences from the same disease but different patients (**Fig. 2C**). We benchmarked our model against ESM2-650M, a general purpose protein language model larger than our model, and Ab-RoBERTA, a recent encoder-decoder antibody protein language model that matches or exceeds the performance of other common antibody protein language models (22). We also compared performance of a similarly trained protein language model and found the DNA language model performed better on the embedding similarity task (**Fig. S2A**). We next used an MLP to classify last-token embeddings of full sequence or junction into healthy, autoimmune disease, infectious disease, other disease, or MS categories. We created two folds in the training data for holdout assessment while preserving grouping by the original study to de-emphasize similarities observed between sequences from the same study. We trained the language model embedding classifier with either a standard cross-entropy loss function or an optimized loss routine designed to emphasize disease-specific predictions (**Fig. 2D**). The optimized loss routine removed MS patient sequences confidently classified to the non-MS condition from the loss function starting at a low rate (e.g. 10% of MS sequences classified to other ignored) before adjusting over training iterations to a higher final rate (e.g. 80% of MS sequences classified to the non-MS condition ignored). We benchmarked the performance of these models against multinomial regression and random forest models trained on nucleotide 7-mer content or an array of protein level physicochemical and sequence features for either the entire VDJ domain or CDR3 alone (**Fig. 2E, F**). Both language models showed superior overall accuracy over baseline models, and the custom loss routine raised prediction precision on holdout test data from a variety of diseases. To test the accuracy of our model in a more stringent context, we compared the classification of MS-derived sequences against only neuromyelitis optica (NMO) patient sequences and found the model distinguished the two diseases via precision in top 5% and 1% of ranked sequences, while performing somewhat worse than when applied to the general disease comparators **(Fig. S2B**). Finally, we analyzed the prevalence of top IGH V genes among classified sequences, finding significant enrichment and de-enrichment of a number of V genes (**Fig. 1G**) and patterns of enrichment among highly used V genes in the dataset (**Fig. 2H**).

**Fig. 2:**
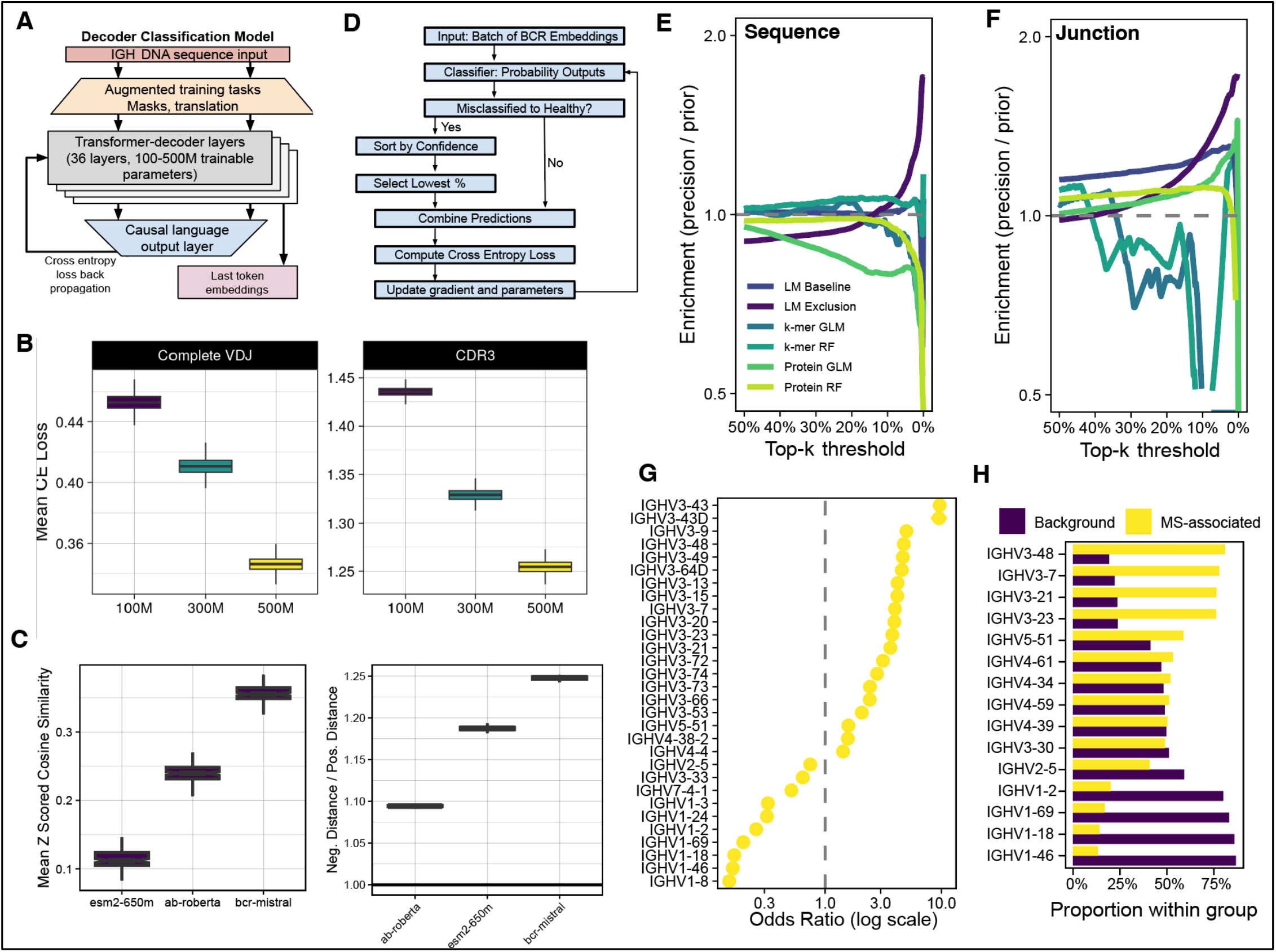
A BCR-specific transformer decoder foundation model and classifier for MS-associated BCRs *A)* Schematic of BCR-Mistral foundation and sequence classifier model architecture, data augmentation, and training process. *B)* Validation set bootstrapped mean cross-entropy loss across complete BCR IGH sequences or BCR IGH CDR3 domains for 500M, 300M, and 100M parameter model sizes. *C)* Comparison of BCR-Mistral embedding performance to Ab-RoBERTa and ESM2-650M by ratio of triplet loss-style positive and negative case differences and mean cosine similarity between antigen-adapted sequences between patients with the same disease. *D)* Schematic loss function optimization procedure that emphasizes high precision for disease-associated sequences. *E)* Test set classification precision across top rank percentiles of various benchmark models and language model classifiers on complete BCR IGH sequences. *F)* Test set classification precision across top rank percentiles of various benchmark models and language model classifiers on BCR IGH CDR3 sequences. *G)* Enrichment of top significantly enriched and de-enriched IGH V genes among MS-classified sequences. *H)* Enrichment of top expressed IGH V genes among MS-classified sequences relative to sequences not classified as MS-associated.

### MS-associated BCRs bind disease-relevant targets and shift distribution in response to therapy

To test if the sequence classifier could be clinically or experimentally useful, we applied the sequence classifier to several patient and antibody stratification tasks. We tested how well top percentile fractions of MS-classified sequences ranked MS patients and found that our dynamic exclusion embedding classifier outperformed the baseline embedding classifier (**Fig. 3A**). To determine if response to treatment could be captured by our simple repertoire-fraction metric, we examined several datasets with patients assayed before and after treatment with natalizumab or fingolimod (**Fig. 3B**). This analysis suggested a decrease in detected disease-associated sequences in Fingolimod patient blood samples (p = 0.12, n = 10) and a decrease in detected disease-associated sequences in high sequencing depth natalizumab-treated patient CSF samples (p = 0.10, n = 4) but not their paired blood samples, although no comparisons reached significance. We observed no trend in detected rate of MS-associated sequences in blood repertoires from a number of other treatments such as ocrelizumab and dimethyl fumarate (**Supp. Fig. 2**). However, many of these samples contained very low sequence sample depth along with small patient counts, likely affecting the power of the analysis. Using blood samples only from patients with the three most common disease state classifications, we calculated a simple metric of the percent of the frequency-weighted repertoire that was classified as MS-associated (**Fig. 3C**). This revealed clear differences in detected disease-associated repertoire fraction, with patients scoring lowest at diagnosis (CIS) and primary progressive patients scoring higher, though formal significance testing was precluded by sampling correlation with study of origin and patient age. To determine if MS-associated sequences encode putative disease-associated antibodies, we classified IGH sequences from four previously described proteolipid protein 1 membrane complex (PLP1c)-binding antibodies retrieved from MS CSF plasmablasts and an isotype control antibody (24). Three of four of the PLP1c-binding sequences ranked above the isotype control antibody along their probabilities of MS-association (**Fig. 3D**). We used junction embeddings for these analyses due to missing nucleotides in some of the complete V-J sequence alignments derived from Sanger sequencing. We validated the specificity of these antibodies and controls via live binding to O4+ oligodendrocytes using flow cytometry (**Fig. 3E**). These results together indicate the utility of our language model-based sequence classification approach for identifying patient disease states and disease-related antibodies.

**Fig. 3:**
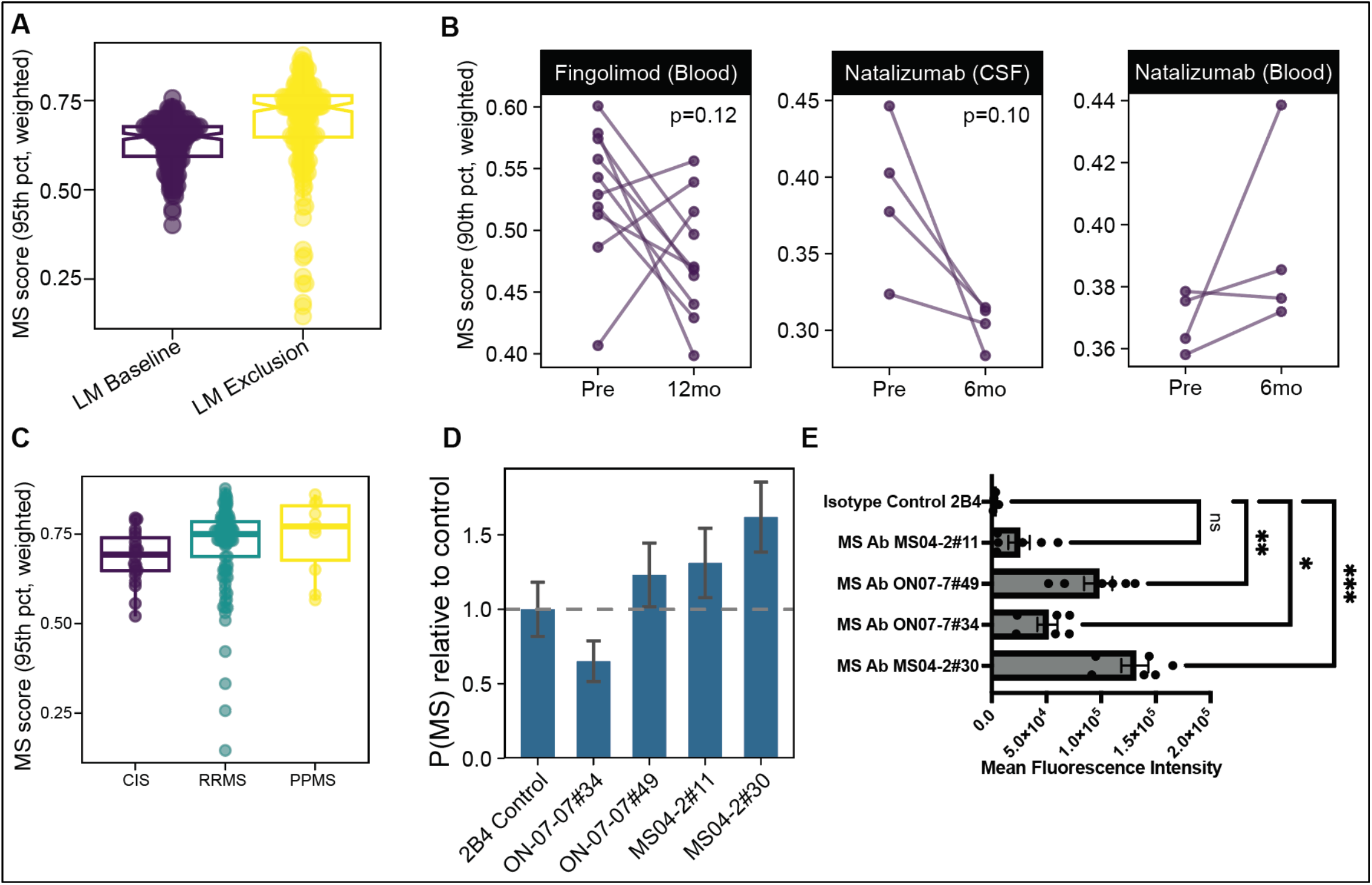
MS-associated BCRs bind disease-relevant targets and shift distribution in response to therapy *A)* Mean weighted probability of MS patient blood repertoires classified as MS-associated in 95^th^ percentile of score-stratified sequences compared between language model embedding classifiers with and without dynamic label exclusion. *B)* Comparison of mean weighted probability of MS patient blood repertoires classified as MS-associated in 90^th^ percentile of score-stratified sequences before and after treatment with Fingolimod (blood samples), Natalizumab (CSF samples), and Natalizumab (blood samples). Paired Mann-Whitney U test. *C)* Mean weighted probability of MS patient blood repertoires classified as MS-associated in 95^th^ percentile of score-stratified sequences compared between MS disease classes (CIS, clinically isolated syndrome; RRMS, relapsing remitting MS; PPMS, primary progressive MS). No significance indicated due to sampling correlation with study of origin and age. *D)* Junction embedding classification MS probabilities of five assayed antibodies cloned from MS patient or control patient CSF plasmablasts normalized to control antibody MS probability. *E)* Mean fluorescence intensity of indicated recombinant antibody binding measured to O4+ adult mouse oligodendrocytes via flow cytometry. Ordinary one-way ANOVA with Dunnett’s post-hoc test. *** p < 0.001, ** p < 0.01, * p < 0.05.

### MS-associated BCRs and BCR repertoire embeddings allow the classification of MS patients from other diseases

Because sequence-disease associations accurately stratified disease-associated sequences and disease states, we hypothesized that MS patients could be accurately distinguished from other diseases using sequence predictions and/or unsupervised embeddings. We first used our simple repertoire fraction score to classify patients (**Fig. 4A**) yielding weak classification (ROC AUC 0.66). To capture variable-sized repertoires into fixed size inputs for patient classification, we defined fixed waypoints in the BCR embedding space and measured the proximity of sequences to each waypoint to yield a 128-dimensional repertoire re-embedding (**Fig. 4A,B**). Waypoint locations were optimized using either a reconstruction objective via a linear decoder or via a joint reconstruction and contrastive unsupervised objective. Waypoint embeddings alone or waypoint embeddings paired with sequence predictions yielded stronger patient classification (**Fig. 4B**). Patient classification performance was only slightly diminished when evaluated in blood repertoires only (**Fig. 4C**). Because many patient repertoires assembled from archival data were based on very few sequences, we enforced maximum thresholds of 100, 200, 500, 1000, or 10,000 sequences per repertoire and found that patient classification improved as repertoire size increased (**Fig. 4E**). The strongest model, a boosted forest model with waypoint embeddings and sequence predictions combined as inputs, performed well across healthy, infectious disease, and autoimmune disease comparator repertoires, with performance falling slightly in the diseased comparisons (**Fig. 4D**). We used SHAP (Shapley additive explanations) analysis to determine which waypoints were significantly associated with increasing or decreasing patient classification to MS. We then found the 1000 sequences nearest to each of these waypoints in the original BCR embedding space and determined which V genes were enriched for each waypoint (**Fig. 4F**). These results show a strong ability of BCR embeddings, even with heterogeneous archival data, to classify MS patients from autoimmune, healthy, and infectious disease controls.

**Fig. 4:**
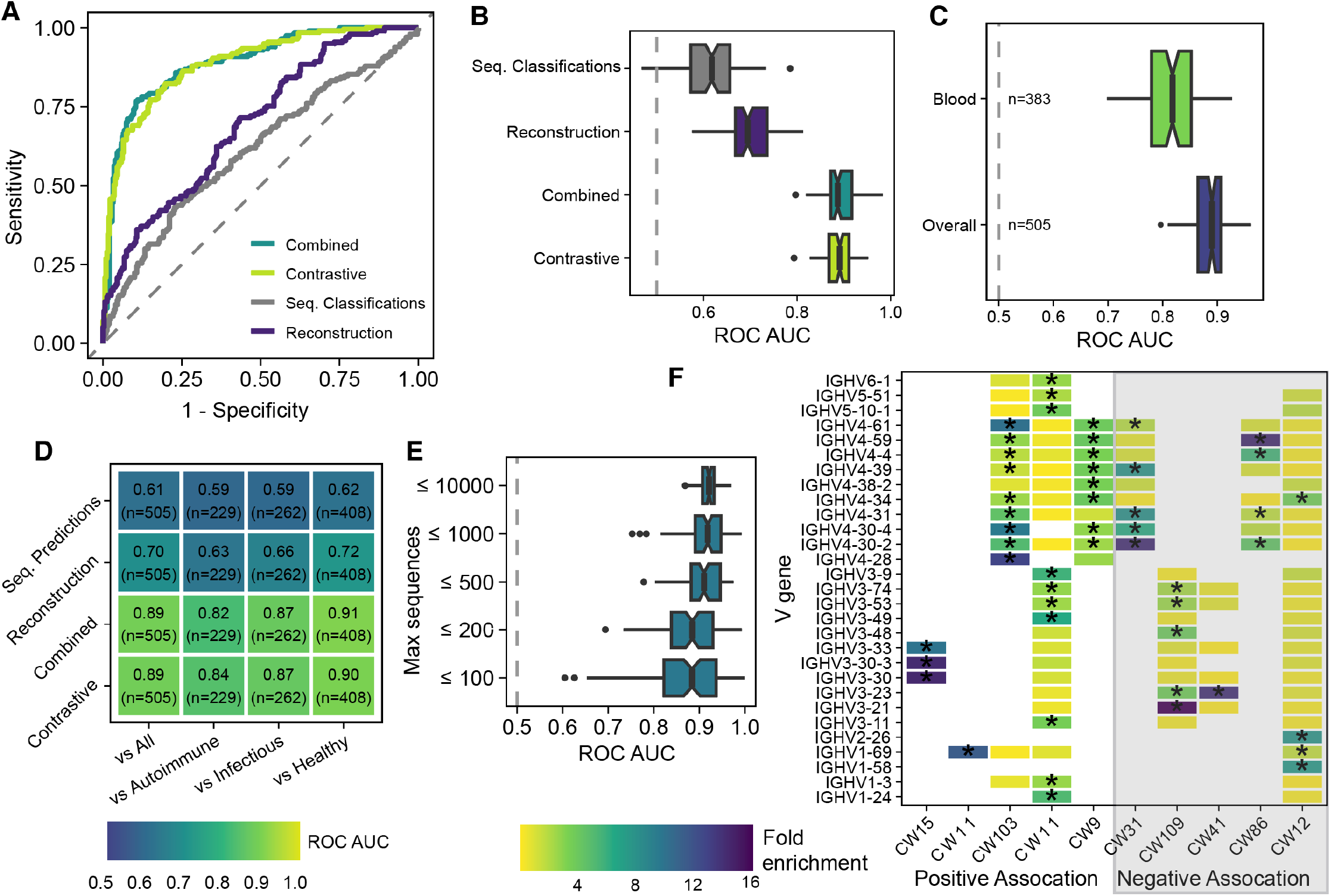
Sequence classifier model associations and BCR embeddings allow classification of MS patients from other diseases *A)* ROC curves showing classification performance for MS vs all sequence classification metrics, reconstruction tuned waypoints, contrastive and reconstruction tuned waypoints, and contrastive and reconstruction tuned waypoints plus sequence classification metrics. *B)* Study level bootstrapped ROC AUC of indicated model data inputs. N=505 repertoires. *C)* Study level bootstrapped ROC AUC of contrastive waypoint model for blood repertoires (N=383) vs all repertoires (N=505). *D)* Heatmap showing performance of contrastive waypoint model using all repertoires above 100 sequences against individual disease categories. *E)* Bootstrapped mean ROC AUC of contrastive waypoint model across various maximum repertoire sizes. N=55 max 100, N=97 max 200, N=153 max 500, N=188 at max 1,000, N=465 repertoires at max 10,000. *F)* V gene enrichment among 1000 sequences nearest to top 5 significant MS-associated waypoints and top 5 other disease associated contrastive waypoints via SHAP analysis. * FDR < 0.001. Wilcoxon rank-sum test.

## Discussion

In this work, we developed and analyzed a unique resource of MS patient BCR sequences, much of which was extracted for the first time from bulk RNA sequencing databases. These findings demonstrate the utility of archival bulk-extracted adaptive immune reads, an underutilized target of data reuse. Although the absolute number of sequences extracted is large, we believe the primary value of this resource is its heterogeneity across sampling methods, geographies, patient states, and sequencing technologies. Compiling this resource required considerable manual curation of metadata, and we will not be the first to note that available metadata in the sequence read archive is critically underreported and unstandardized. Using these curated data, we were not only able to recapitulate and extend findings about V gene usage and CDR3 length in MS but also to show that many putative public clones are consistent with technical inflation from sample preparation or index hopping, though contributions from shared patient characteristics within studies cannot be excluded. This observation underscores the need for BCR analysis tools that capture the higher-order features of each sequence to understand their relationship to disease.

We built a BCR-specific DNA-language model to better extract features related to disease among the sequences in our dataset. We turned to language model-based sequence embedding after being disappointed by the performance of several more traditional DNA and protein feature approaches that did not yield sufficiently predictive models. Our model choice is in several ways unorthodox. First, we chose to train our model at the level of DNA nucleotides in order to capture known MS-relevant features like codon bias and silent mutations while also capturing all information available to a protein language model (25, 26). Second, we chose to derive our architecture from a decoder-only model to create a parameter-efficient embedding model without dropping the decoder after pretraining. Finally, at 500 million trainable parameters, our model is considerably larger than prior BCR protein language models. Despite the increase in parameters, we were able to achieve superior embedding performance with a limited amount of pretraining resources by creating a highly diverse training dataset using a highly efficient sampling method. Our methods created expressive embeddings that outperformed both a state-of-the-art BCR embedding model and a state-of-the-art general-purpose protein language model that has more parameters than our model. Expanding this approach to other adaptive immune receptor chains and other genomic embedding tasks should prove valuable.

In developing the methods presented here, we frequently confronted the subtle and confounding ability of machine learning models, especially very powerful ones, to capture irrelevant but predictive features of the input data. We found that the BCR assembly method and other study-specific variables, like the sequencing library preparation method, were easily detected by our models; removing these signals required considerable regularization steps. Another source of bias that we highlighted is public clones, which may be technical artifacts. This conclusion is supported by the much higher sequence classification performance we observed when we created train-test folds for the sequence classifier based on patients rather than studies.

Beyond these splits, our custom loss routine is designed to focus a classifier on the small proportion of disease-relevant signals in a BCR repertoire that can easily be otherwise overwhelmed by technical signals. By removing sequences confidently classified to the non-MS class during training but retaining a fraction of such misclassifications, we create a training regimen analogous to semi-hard example mining. Similarly, at the patient level, we observed clustering by study and poor patient classification when we used cross-validation that preserved study grouping without batch correction. True out-of-study validation is somewhat uncommon, and we would like to highlight how our design across many heterogeneous studies increases the validity of the analysis.

The utility of the approach developed here is demonstrated by our analyses of patients before and after treatment and classification of PLP1c-binding antibodies. We found that, across studies, pre- and post-treatment sample data were variable; nevertheless, the impact of fingolimod and natalizumab on blood and CSF repertoires are consistent with their effects on circulating/migrating B cell populations. Large sample sizes will be required to measure treatment-related repertoire changes accurately. An important future application of these metrics could include monitoring treatment response and tolerization. We demonstrated that our model can correctly classify a small sample of myelin protein-binding antibodies. Considerable interest in the field has been directed towards identifying epitopes targeted in MS, and we believe that sequence classification tools like the ones described here could be productively applied to identifying and testing candidate sequences.

Several important questions remain unaddressed by our work. It is reasonable to assume, and we expect, that TCRs and immunoglobulin light chains contain relevant information about MS disease state. Incorporating these receptors and capturing the relationship between them with each patient will likely be important to raising our patient-level predictive accuracy to clinically useful levels. Although our archival data contain a wide variety of neurological and other conditions, and we used autoimmune conditions in the OPIG OAS as comparators, more data from genuinely clinically relevant comparators like neuromyelitis optica (NMO), myelin oligodendrocyte glycoprotein antibody-associated disease (MOGAD), and other autoimmune neurological conditions will be needed in the future, both to develop strong models and to test their usefulness. Similarly, our archival approach yielded highly undersampled repertoires for many patients. Future efforts to collect large cohorts of deep and standardized repertoires will be needed to further the field. In summary, we present a dataset, foundation model, sequence, and patient classifiers that should prove useful towards detecting and analyzing MS disease states, as well as other autoimmune and infectious disease states.

## Materials and Methods

### Systematic Data Collection

Archival data from studies containing patients with multiple sclerosis were identified in the NCBI SRA archive using the search terms: (multiple sclerosis) AND “Homo sapiens”[orgn: txid9606]. Data were filtered to include only mRNA sequencing data (4,449 files filtered), remove metagenomic samples (383 files removed), and to remove sequencing from purified non-immune cell types (e.g. purified oligodendrocytes, thrombocytes), 3’ selected 10X Genomics single cell data (which does not typically capture CDRs), and to remove duplicated samples in the archive (4 duplicated study submissions identified). Metadata was added where available from manuscripts and supplemental information for the original studies. Full metadata for SRA runs included and filtered is included in the data archive associated with this work. Additional BCR IGH data were collected from the OPIG OAS unpaired BCR archive after filtering for human samples that contain individual patient IDs and deduplicating studies found in both SRA and OPIG OAS (17).

### TRUST4 Adaptive Immune Receptor Assembly

Data identified from SRA were accessed using the SRA toolkit package (v3.0.0) and converted to paired or unpaired fastq files as appropriate. Files were trimmed and quality filtered using fastp with default settings, then candidate BCR sequences were extracted and assembled using TRUST4 (16, 27). Files that failed to complete assembly in the first pass were re-run using the --repseq setting to reduce the computational cost of alignment. Pyrosequencing files were subjected to more strict quality filtration settings. Single-cell files were assembled using appropriate cell- and molecule-level barcode demultiplexing. Sequences from light chain and TCR loci, with any ambiguous bases, a stop codon, and those did not contain the V gene start through the J gene end were filtered from the dataset. Extracted sequences were re-annotated using IgBLAST to ensure the consistency of germline gene assignment and V gene start and J gene end calls with the OPIG OAS data workflow, resulting in the removal of an additional 19.2% of sequences that did not meet the filtration criteria above after re-annotation (28).

### Transformer-Decoder BCR DNA Foundation Model

Three sizes of decoder-only transformer-based language foundation models were pre-trained on next-token prediction tasks (23). We trained a large model with approximately 500 million trainable parameters, a medium model with 300 million trainable parameters, and a small model with 100 million trainable parameters. For pretraining data, OPIG OAS was filtered to include only IGH sequences, then transformed into 7-mers. Sequences were sampled based on their Euclidean distance in 7-mer space from an iterative random sample of sequences to achieve a compact (∼2 million sequences) training dataset of highly diverse sequences. During pretraining, sequences were tokenized at the nucleotide level, randomly rotated, masked with up to two 20-nucleotide attention masks, and randomly truncated up to a third of their length after rotation. The foundation model variants were trained using ROCm 5.2.3 on AMD MI100 GPUs on the RMACC Alpine supercomputer. Cross-entropy loss was minimized via the AdamW optimizer with weight decay of 1e-2 after gradient accumulation to an effective batch size of 2400 sequences (approximately 320,000 nucleotide predictions). Learning rate was annealed from 1e-5 to 1e-7 across pretraining. Each model was trained for an equivalent amount of clock time to approximate similar computational resources. After comparing model sizes, the large model was used for all subsequent tasks. A similar foundation model was trained on junction sequences only and used for the junction embedding classifier inputs.

### BCR Embedding Sequence Classifiers

Language model sequence embeddings were derived from the large unsupervised foundation model. We then used a multilayer perceptron with three layers of size 256, 128, and 64 to classify these embeddings. For classifier training, patient disease labels were propagated to each sequence and then assigned to either healthy, autoimmune disease, infectious disease, other disease, or multiple sclerosis categories. A maximum of 20,000 BCR IGH sequences with complete VDJ regions and predicted productive status were randomly sampled from each patient in the dataset. Data were split into two equal-sized folds with samples from the same studies grouped together such that they were all assigned to one or the other fold. Cross-entropy loss was used to optimize for default models with cosine annealing of the tuned learning rate after a warmup period. Models were tuned across three inner folds grouped by study within each outer fold before being used to predict as an ensemble of 10 models on the outer validation fold. A batch size of 2048 was used for all models. The domain-specific loss function for the optimized model dynamically excludes a fraction of sequences from the multiple sclerosis category that are classified to the non-MS label for the purposes of loss calculation in each training batch. The excluded sequences are selected from the most confident healthy misclassifications. The fraction excluded was tuned between a low starting point (0-25%) and a final higher rate (25-100%) along with the step size change in exclusion rate and over training using a nested cross validation strategy within each training fold.

### Live Cell Antibody Binding Assay

For flow cytometric analysis of recombinant human antibody binding, primary cells were dissociated from adult mouse brains using a collagenase-based dissociation (Miltenyi), blocked with normal donkey serum (NDS), then incubated with 1 µg recombinant antibody per 100,000 cells for 1 hour at 4°C. Cells were washed and then incubated with donkey anti-Human IgG AF594 and anti-O4-APC. Per-cell fluorescence among O4+ cells was measured on a Cytoflex LS flow cytometer. Cytometry data were analyzed using Floreada software. Fold change was calculated between antibody binding to O4+ and O4-cells within each sample.

### Waypoint-Based Repertoire Embedding

For waypoint-based patient embeddings, sequences were passed through the large foundation model, and hidden states corresponding to the last nucleotide token were extracted from the final layer of the model. 128 waypoints were randomly initialized in the hidden state space. The Euclidean distance between a sequence and each waypoint was measured and normalized as 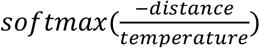. Repertoires were summarized as the mean scaled distance between each waypoint and all sequences in the repertoire. These waypoints were then optimized in the hidden-state space using reconstruction mean squared error loss after passing the normalized waypoint distances through a single-layer linear decoder over the training dataset for the “reconstruction” waypoints and with a joint decoder reconstruction and contrastive temperature normalize cross-entropy loss minimization for the “contrastive” waypoints. For contrastive error, unlabeled repertoires were compared to sampled subsets of the same repertoire and the distance between subsets was minimized while maximizing the distance from the comparator repertoire. Final waypoint locations were fit across all repertories. For inputs to the patient classification model, patients were summarized as the mean normalized distance from each waypoint to every sequence in the BCR repertoire.

### Patient Classification Models

Patient classification models used summarized sequence classification predictions and/or waypoint-based repertoire embeddings. Sequence classifications were made by two models on their respective cross-validation test sets. Waypoints were optimized as described above using unsupervised reconstruction error. Waypoint re-embeddings were then calculated for all repertoires in the dataset. Waypoint re-embeddings were batch-normalized for tissue source before patient classification using Limma (29). For classification using the fraction of MS-associated repertoire alone, we fit simple logistic regression models. For patient classification using waypoints or waypoints with MS-associated repertoire fraction, we fit gradient boosted forest (XGBoost) models using 500 trees across 5-fold cross-validation with validation folds split by study and grouped by studies from the same lab. Model hyperparameters were tuned within the four inner folds before predicting on the outer validation fold. SHAP analysis was implemented through the xgboost package in R (30). Significant SHAP values were selected by FDR < 0.05 across patient resamples.

### Classification Comparator Models

Logistic regression and random forest comparator models were trained on nucleotide sequence and protein feature data. Models were trained on randomly downsampled 200,000 sequence subsets of the combined TRUST4 and OPIG OAS dataset to maintain reasonable runtime. Sequence models used nucleotide 7-mers filtered for zero variance features. Protein feature models used pseudo amino acid composition and Moran autocorrelation implemented in the protr package with highly correlated features removed prior to training (31). Logistic regression was fit as lasso regression with L1 regularization of 0.01. Random forest models were trained with 500 trees and a minimum node size of 5. All models used case weighting scaled to tissue source as described above. Models were evaluated using the same 2-fold cross validation folds as the language model embedding classifiers.

### Recombinant Antibodies

Immunoglobulin sequences were cloned from patient intrathecal B cell repertoires and expressed as full length human IgG1 as previously described (32, 33). Briefly, bivalent human IgG1 antibodies were expressed in EXPI293 cells (ThermoFisher) and purified with protein A-Sepharose (Sigma-Aldrich). The PLP1c binding clones MS04-2#30, MS04-2#11, ON07-7#34, and ON-07-7#49 were used for all studies along with indicated isotype controls (24).

### V Gene and Public Clone Analyses

V gene analyses among classified sequences (Fig. 2G, 4F) use Fisher exact tests of proportions between classified or non-classified sequences or associated and background sequences. For V gene frequency analyses in Fig. 1F and G, frequencies were calculated per repertoire in samples with more than 100 clones and compared using a Kruskal-Wallis test with FDR adjusted p values. For the public clone analysis, we deduplicated clones at the CDR3 amino acid level for patients or studies, depending on the contrast. Patients without unique identification within their study were excluded. We estimated an empirical rate of public clones across all patients across all studies at various sample sizes as a baseline. This baseline was used to estimate expected CDR3 collisions based on sample sizes between patients and studies. We then sampled patients within studies or across studies and determine the relative enrichment of CDR3 collisions compared to this baseline rate. Patients and studies were bootstrap resampled 100 times to estimate variance.

## Data and Code Availability

Code used for analyses in this paper will be available upon publication at https://github.com/krishnanlab/bcr-mistral-final. Raw and processed datasets used in this paper along with model weights will be made available at https://doi.org/10.5281/zenodo.20432913.

## Acknowledgements

This work utilized the Alpine high-performance computing resource at the University of Colorado Boulder. Alpine is jointly funded by the University of Colorado Boulder, the University of Colorado Anschutz, Colorado State University, and the National Science Foundation Award #2201538. Claude Opus 4.6 was used for some coding tasks associated with this work. This work was supported by NINDS F31NS141369 to G.C.P., NIGMS R35GM128765 to A.K., and NINDS R21NS142515 to W.B.M.

## Abbreviations

MS: Multiple sclerosis
CSF: Cerebrospinal fluid
SRA: Sequence read archive
BCR: B cell receptor
TCR: T cell receptor
NMO: Neuromyelitis optica
IGH: Immunoglobulin heavy chain

---

**Supplemental Figure 1:**
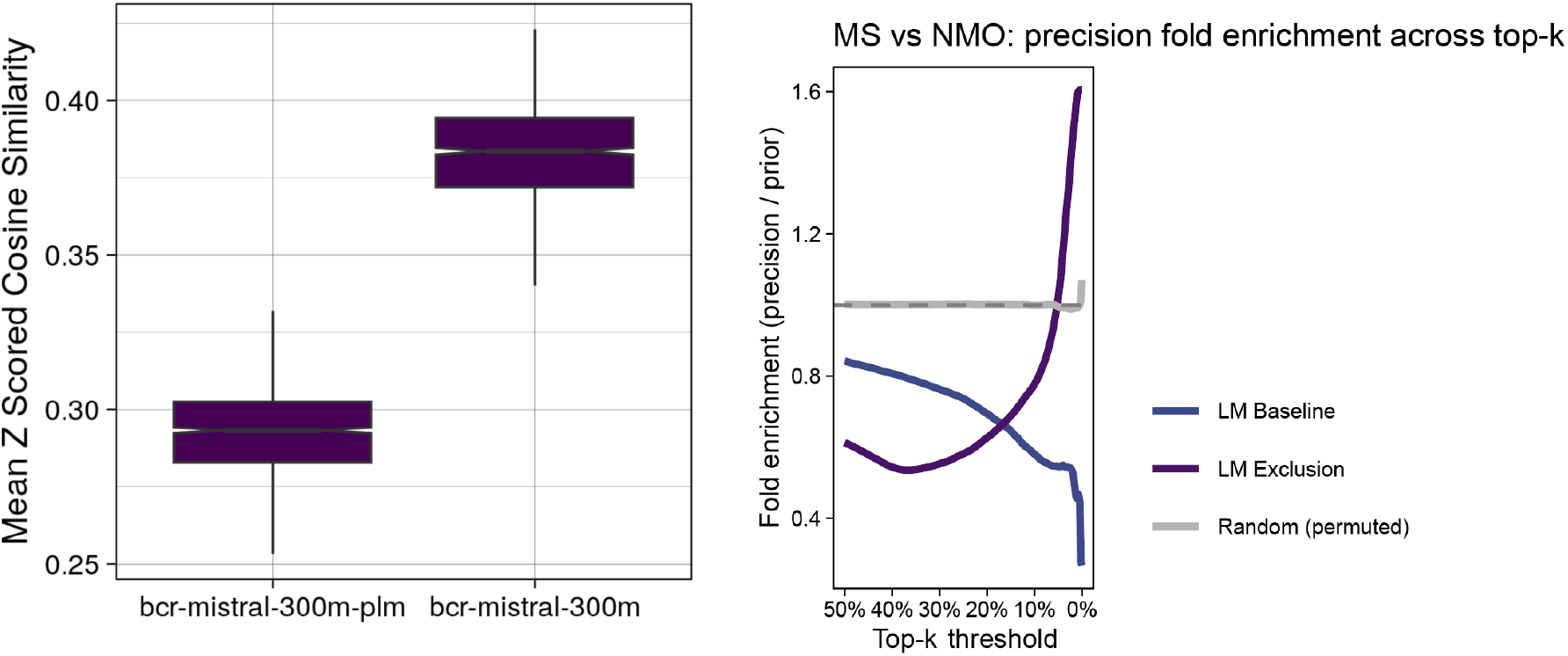
Protein BCR-Mistral variant and MS vs NMO Precision

**Supplemental Figure 2:**
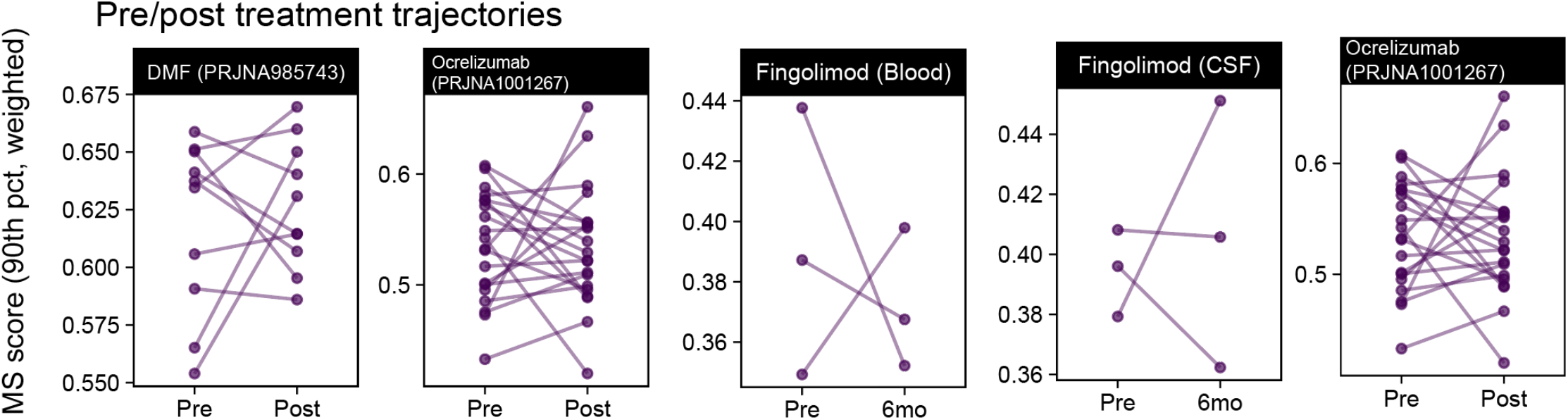
Additional pre-post treatment statistics

